# Population analysis and immunologic landscape of melanoma in people living with HIV indicate impaired antitumor immunity

**DOI:** 10.1101/2025.04.17.648995

**Authors:** Lindsay N Barger, Derek Wang, Ashley L Saravia, Valeria Mezzano, Gyles Ward, Cynthia Loomis, Carly Feldman, Madalina Tuluc, Rino S Seedor, Peter J Gaskill, Anna E Coghill, Gita Suneja, Iman Dehzangi, Jennifer L Hope, George Jour, Gabriele Romano

## Abstract

**Purpose:** To dissect the clinical and immunological features of people living with HIV (PLWH) diagnosed with melanoma, who have consistently shown worse outcomes than HIV-negative individuals (PLw/oH) with the same cancer.

**Experimental Design:** We analyzed electronic health records from 1,087 PLWH and 394,437 PLw/oH with melanoma. Demographic and clinical characteristics were compared. Spatial immune transcriptomics (72 immune-related genes) was performed on melanoma tumor samples (n=11), with downstream validation using multiplex immunofluorescence (n=15 PLWH, n=14 PLw/oH).

**Results:** PLWH were diagnosed at a younger age, had greater representation of Hispanic and Black individuals, and showed reduced survival. They also had a markedly increased risk of brain metastases. PLWH experienced significant delays in initiating immune checkpoint inhibitor (ICI) therapy and had worse post-ICI survival, even after balancing covariates. Spatial transcriptomics revealed a more immunosuppressive tumor microenvironment in PLWH, with increased transcription of immune checkpoints (PD1, LAG3) and reduced antigen-presentation markers (HLA-DRB, B2M), with distinct spatial distributions in tumors and surrounding microenvironments. Multiplex immunofluorescence demonstrated features of an exhausted CD8⁺ T cell compartment, including enrichment of PD1^int^LAG3⁻ and PD1^int^LAG3⁺ subpopulations, and a significant accumulation of myeloid-derived suppressor cells (CD11b⁺ HLA-DR⁻ CD33⁺).

**Conclusions:** Melanoma in PLWH is associated with distinct clinical and immunological features, including delayed ICI treatment, reduced survival, and an immunosuppressive microenvironment with exhausted CD8⁺ T cells and expanded myeloid-derived suppressor cells. These findings suggest that chronic HIV infection may impair antitumor immunity in melanoma. Targeting the pathways identified here may improve therapeutic responses and outcomes in this population.

**Statement of translational relevance:** This study reveals critical barriers to effective melanoma treatment in people living with HIV (PLWH). Despite receiving immune checkpoint inhibitors (ICIs), PLWH face delayed therapy initiation, a greater likelihood of brain metastases, and significantly higher long-term mortality, even after adjusting for demographic covariates. Transcriptional immune profiling further uncovers a tumor microenvironment enriched in immunosuppressive myeloid-derived suppressor cells and CD8⁺ T cell populations with features of exhaustion. These findings suggest that poorer outcomes in PLWH stem not only from delayed care, but also from distinct targetable mechanisms of immune dysfunction. For example, strategies to reverse MDSC accumulation in the tumor or tailored ICI regimens could enhance immune responsiveness and improve treatment efficiency. By defining the clinical and immunological features of this population, this work highlights opportunities for precision immunotherapy tailored to PLWH with melanoma, with direct implications for improving survival and reducing disparities.

## Introduction

Advancements in antiretroviral therapies (ART) have transformed HIV infection from a terminal diagnosis into a chronic condition, allowing people living with HIV (PLWH) to achieve a life expectancy nearly equal to that of the general population ^1,2^. However, despite the success of ART in suppressing viral replication, HIV remains a lifelong infection due to its ability to integrate into host genomes and establish a state of latency, particularly within long-lived memory T cells and myeloid cells in sanctuary sites ^3–5^. While ART prevents the spread of the infection, it does not eliminate pre-existing reservoirs that harbor integrated and latent viruses, leaving PLWH with persistent immune dysregulation and, therefore, increased susceptibility to age-related diseases ^6^. Among these, non-AIDS-defining cancers (NADCs) have emerged as a major cause of mortality, including cutaneous melanoma ^6^. Several reports on melanoma in PLWH point to disproportionately worse clinical outcomes compared to melanoma patients without HIV, despite adjustment for key factors such as disease stage at presentation ^7–9^. The mechanisms underlying this disparity remain poorly understood. It is largely unknown how chronic HIV infection alters anti-tumor immune responses, which is particularly relevant in melanoma, where immune regulation is central to disease progression and treatment response.

To address this major knowledge gap, we first analyzed publicly available clinical datasets to compare demographics and outcomes between PLWH and people living without HIV (PLw/oH), revealing significant disparities in demographics, metastasis risk, and survival after a melanoma diagnosis. We then performed the first immune-focused spatial transcriptomic analysis of melanoma tumors according to patient HIV status, uncovering striking differences in tumor-associated T cell and myeloid populations in PLWH. These findings were further validated and expanded using multiplex immunofluorescence to characterize immune cell phenotypes and subpopulations. Together, our data reveal a profoundly altered immune landscape in melanoma among PLWH, marked by T cell exhaustion and myeloid-mediated suppression, which underscores the need for tailored therapeutic strategies in this population with suboptimal clinical outcomes.

## Methods

### Clinical electronic record patient dataset

The data used in this study were collected on 10/08/2025 from the TriNetX Research Network, which provided access to electronic medical records (diagnoses, procedures, medications) from approximately 157,574,584 patients from 111 healthcare organizations. Patient demographics include date of birth, age, sex, marital status, race, and ethnicity. Demographics are coded to HL7 version 3 administrative standards. For the survival studies, the date a patient was reported as “Deceased” in the dataset was used; the cause of death was not available in the dataset. The “Deceased” term in the TriNetX data model pools together several different sources of mortality data: documented mortality by HCOs using ICD-10 code R99 (ill-defined and unknown cause of mortality); if mortality is reported from HCOs via death dates of patients, this is also captured and mapped to the “Deceased” term; for the subset of patients on the TriNetX networks that have additional mortality data linked to their electronic health records (closed claims, social security master death file, private obituaries, etc.), these additional sources are also mapped to the “Deceased” term. Diagnoses and medication data in the TriNetX database are obtained from electronic health records of participating healthcare organizations. Diagnoses are recorded using the ICD-10 standardized coding system, while medication information is captured through prescription and dispensing records. These data may originate from clinical documentation, problem lists, billing records, or insurance claims and are harmonized across sites to ensure consistency.

Two cohorts of patients were retrieved with the following criteria: PLWH cohort was built selecting patients that had Malignant melanoma of skin (ICD code C43) and Human immunodeficiency virus [HIV] disease (ICD code B20, 1,087 patients were retrieved); PLw/oH cohort was built selecting patients that had C43 but not code B20 (394,437 patients were retrieved). Only patients with a cancer diagnosis between 2005 and 2025 were included. Only patients with a cancer diagnosis after HIV were included in the PLWH cohort.

The following ICD codes were used for metastatic disease: C78.7 (liver and bile duct), C79.3 (brain and meninges), C77 (lymph nodes), and C78.0 (lung). Two different analyses were performed for patients with an early occurrence of metastatic disease (from day 1 to 6 months after diagnosis) and patients with a later occurrence of metastasis (6 months from melanoma diagnosis onwards). For the “All sites” category, patients who had any of the metastatic codes C77, C78.0, C79.3, and C78.7 were included. For co-morbidity assessment, the following codes were used: I20-I25 (Ischemic heart diseases), I11 (Hypertensive heart disease), J40-J4A (Chronic lower respiratory diseases), N18 (Chronic kidney disease CKD), B18 (Chronic viral hepatitis), and E10 (Type 1 diabetes mellitus). In the dataset, medications are represented at the level of ingredients, coded to RxNorm, and organized by VA or ATC classification. For the analysis of immunotherapy characteristics and outcomes, the two cohorts were filtered to include only patients who had received any of the following FDA-approved immune checkpoint inhibitors: 1094833 (ipilimumab), 1597876 (nivolumab),1547545 (pembrolizumab), 1792776 (atezolizumab), 1919503 (durvalumab), or 2596773 (relatlimab).

### Inclusion and Ethics

These studies were conducted in accordance with recognized ethical standards, including the U.S. Common Rule and the Declaration of Helsinki. The retrospective study was exempt from informed consent and Western Institutional Review Board (IRB) review. All data analyzed were part of a secondary analysis of existing records, involved no intervention or interaction with human subjects, and were fully de-identified in accordance with the de-identification standard outlined in Section §164.514(a) of the HIPAA Privacy Rule. De-identification was performed and formally certified by a qualified expert, as defined in Section §164.514(b)(1), with the determination most recently renewed in December 2020.

### Spatial Immune Transcriptomics

De-identified melanoma patient samples were acquired from the AIDS and Cancer Specimen Resource, the Sidney Kimmel Comprehensive Cancer Center, and NYU Grossman School of Medicine. Sample collection and handling were evaluated by institutional IRB. 11 melanoma samples from people living with HIV were retrieved first, and 11 HIV-uninfected controls were then selected to match age, sex, race, and ethnicity, and nature of specimens (**Table S1**). Five-micron sections were prepared after cleaning the microtome and recovering sections in RNAse-free water. Slides were processed per NanoString® protocols for nCounter RNA GeoMx experiments. Slides were baked at 60°C for 2 hours and loaded onto a BOND Rx stainer (Leica). Antigen retrieval was performed with EDTA-based solution (pH 9, ER2 Leica, AR9640) at 100°C for 20 minutes, followed by proteinase K (Ambion, AM2546) treatment (1 μg/mL, 15 minutes, room temp). Slides were stored in PBS and incubated overnight with 200 µL of GeoMx Immune Pathways Hs RNA Probes (121300201) at 37°C in a HybEZ™ oven under a Hybrislip. Post-hybridization, slides underwent two washes in 50% Formamide-4XSSC at 37°C and two washes in 2XSSC at room temperature, then labeled with RNA-compatible melanoma morphology markers (GeoMx Melanoma Morph Kit Hs RNA). All slides were processed together but loaded onto the GeoMx instrument in batches of four. Automated fluorescence imaging at 20X magnification (0.4 μm/px) was followed by ROI selection. For each sample, three to five regions of interest (ROIs) were selected for analysis. ROI selection and masking were performed in collaboration with a board-certified dermatopathologist to ensure accurate discrimination between tumor and tumor-microenvironment compartments. ROIs were identified based on morphological assessment and multiplex immunofluorescence staining for S100B/Pmel17 (melanocytic markers), CD45 (pan-leukocyte marker), and DAPI (nuclear counterstain). Tumor-cell–rich (TUMOR) regions were defined as areas enriched in melanoma cells, characterized by cohesive nests of S100B/Pmel17⁺ cells with consistent nuclear morphology and limited CD45⁺ immune infiltration. Tumor-microenvironment–rich (TME) regions were defined as areas negative for melanocytic markers (S100B/Pmel17⁻) and enriched in CD45⁺ immune cells and/or stromal components, representing the surrounding immune and stromal microenvironment. Following ROI identification, digital masks were applied to precisely capture TUMOR (green) and TME (red) subregions (**Fig. S1**). ROIs were UV-illuminated (∼1 μm² resolution) to release barcodes, which were aspirated and transferred to a 96-well plate. Eluted material was stored at -20°C until processing at NYUGSM. Plates were lyophilized at 65°C for 1–2 hours, rehydrated in DEPC water, mixed with ICP-hyb buffer and GeoMx Hyb Code Pack_RNA probes (121300402), and incubated overnight at 65°C. Hybridized samples were pooled by column and run on the nCounter MAX/Flex system to generate RCC files, which were analyzed using the GeoMx Control Center.

Read counts were normalized to the geometric mean of five housekeeping genes (OAZ1, POLR2A, RAB7A, SDHA, UBB). Normalized read counts are reported in **Table S2**. Differential expression analysis (DEGs) was conducted on the normalized read counts using the DEseq2 algorithm ^10^ in R v4.4.2 to compare the TME and TUMOR across HIV-positive and HIV-negative patients.

### Immune pathways and population analysis

The MCP-counter algorithm was used to calculate the enrichment scores of tailored gene sets^11^. We conducted two marker gene–based analyses: one focusing on differences in immune signaling gene expression (53 genes, **Table S3**), and another on estimating immune cell population abundances (19 genes, **Table S4**). A multiple Welch t-test was then performed on the population or pathway scores in HIV+ vs. HIV-samples. Data were represented in a bubble plot with the adjusted p-value (q-value) and Log2 fold change (HIV+ vs. HIV-) for the TUMOR and TME areas (R v4.4.2; ggplot2 package).

### Multiplex Immunofluorescence (mIF)

mIF staining was performed on 5 μm paraffin sections with Akoya Biosciences® Opal™ reagents on a Leica BondRx autostainer, according to the manufacturer’s instructions. In brief, sections were treated with hydrogen peroxide to block endogenous peroxidases, followed by antigen retrieval using ER2 (Leica AR9640) at 100°C. Primary and secondary antibodies were applied sequentially, with each pair followed by HRP-mediated tyramide signal amplification using a unique Opal® fluorophore. After each staining round, antibodies were stripped via heat retrieval, leaving the fluorophore covalently bound at the antigen site. Sections were counterstained with spectral DAPI (Akoya FP1490) and mounted with ProLong Gold Antifade (ThermoFisher P36935). Imaging was performed using a Leica AT2 (40X) or Akoya Vectra Polaris (20X), and qptiff scans were generated with PhenoImagerHT 2.0, Phenochart 2.0, and InForm 3.0. Scans were analyzed in QuPath with four ROIs per sample—two central and two peripheral—independently selected by two operators using fixed-threshold detection. Averages of ROIs and operators were used as replicates. Total cells were detected via QuPath’s DAPI-based algorithm. T cell and myeloid infiltrates were assessed using two antibody panels: Panel 1 (CD8, CD4, FoxP3, Lag3, PD1, **Table S5**) and Panel 2 (CD11b, CD33, HLA-DR, CD66b, **Table S6**).

A custom QuPath script was developed to classify CD8+ T cells and respective PD1 expression into three tiers (low, intermediate, and high), with an additional subdivision for LAG3 positivity. PD1 intensity thresholds were manually defined after visual review by two independent operators to capture biologically distinct expression levels. Based on this consensus, cells with PD1 intensity between 0.0 and 0.2 were classified as PD1 negative, while those with intensities above 3.0 were designated as PD1 high. The intermediate tier (0.2–3.0) captured cells with PD1 expression levels falling between these two extremes. A similar script was generated for CD4+ T cells and relevant subpopulations. CD11b+ myeloid cells and their co-expression with HLA-DR, CD33, or CD66b were quantified similarly.

### Statistical analysis

Data was checked for normality (Kolmogorov-Smirnov, Shapiro-Wilk test) prior to being assigned to parametric or non-parametric analyses. All the datasets analyzed in this work displayed a normal distribution. To compare significant differences between two groups, we used a two-tailed t-test (parametric; Welch correction was applied if variances were different between groups). A p<0.05 was considered significant. For demographic and clinical covariates, univariate T-test analysis was performed. For the immune pathways and population analysis, a multiple Welch t-test was performed with an FDR of 0.01. An adjusted p-value (q-value)<0.01 was considered significant. For survival studies, a Log-rank (Mantel-Cox) test was used to detect differences in survival between two populations. A p<0.05 was considered significant. Hazard ratios (HRs) with 95% confidence intervals (CIs) were calculated using Cox proportional hazards regression. The HR compares the hazard rates—the instantaneous rate at which patients experience the outcome for the first time—between two cohorts. An HR of 1 indicates equal rates of the outcome at any point in time, whereas an HR >1 (or <1) indicates that one cohort experiences the event at a higher (or lower) rate than the other. HR estimates are reported with 95% CIs, and if the CI includes the null value of 1, the difference is not considered statistically significant. The proportional hazards assumption was assessed via the Grambsch–Therneau test based on scaled Schoenfeld residuals, which evaluates whether covariate effects vary over time (a p<0.05 indicates a non-proportional hazard). For risk Ratio assessment, risk ratios with 95% confidence intervals are calculated to compare outcomes between cohorts. Risk was defined as the percentage of patients in the cohort who had the outcome in the time window (Risk = Patients with Outcome/Patients in Cohort). Risk Ratio was defined as the ratio of risk between cohorts (Risk Ratio = Risk for Cohort 1 / Risk for Cohort 2). Cohort balancing in TriNetX was conducted using propensity score–based methods. Propensity scores were estimated via logistic regression models incorporating baseline demographic and clinical covariates. Patients were matched 1:1 using greedy nearest-neighbor matching without replacement, applying a caliper of 0.1 pooled standard deviations of the logit of the propensity score to minimize residual confounding. In addition to matching, inverse probability of treatment weighting (IPTW) was applied as a sensitivity approach, assigning weights based on the inverse of the estimated propensity score to create a pseudo-population in which covariate distributions were independent of exposure status. Balance across matched or weighted cohorts was evaluated by comparing standardized mean differences (SMDs) for each covariate, with thresholds of <0.1 considered evidence of acceptable balance. TriNetX Live was utilized to perform all the population analyses. To ensure patient confidentiality, TriNetX applies an automated privacy protection rule that rounds or suppresses cell counts ≤10. This masking affects only the display of data and does not influence statistical analyses, cohort matching, or aggregate calculations, which are performed using the complete, unmasked dataset. GraphPad Prism v.10.4.1 was utilized to perform all the other statistical tests.

### Data Availability

Raw patient-level data from the TriNetX dataset cannot be publicly shared due to data use agreements and privacy protections. Interested investigators may reproduce our analyses or request access to limited datasets through TriNetX under institutional data-use agreements and IRB approval by contacting TriNetX directly. All analyses were conducted using the platform’s graphical cohort builder and built-in analytics tools; no custom code was used. The complete results of the GeoMX spatial transcriptomics experiment, including ROI-level metadata and normalized expression values, are provided in **Table S2**. mIF raw images are available upon request.

## Results

### PLWH with melanoma display increased overall mortality rates and brain metastasis risk

To evaluate population differences between people living with and without HIV (PLWH and PLw/oH) diagnosed with melanoma, we interrogated the publicly available TriNetX Research Network dataset. Our study included two cohorts: 1,087 PLWH with melanoma and 394,437 melanoma patients without HIV. Univariate analysis of baseline demographics revealed that PLWH were diagnosed with melanoma at a younger age (58.6 ± 12.2 years in PLWH vs. 62.8 ± 15.6 years in PLw/oH) and were more likely to be male (81% vs. 52%, **Table 1**). Racial and ethnic distribution also differed between the total cohorts. PLWH with melanoma exhibited a higher representation of Hispanic or Latino individuals (4% vs. 2%) and Black or African American individuals (7% vs. 1%) compared to their non-HIV counterparts (**Table 1**).

**Table 1.** Demographics of PLWH and PLw/oH with melanoma cohorts for overall survival calculations, before and after balancing cohorts.

|  | Before balancing cohorts |  |  |  | After balancing cohorts |  |  |  |
| --- | --- | --- | --- | --- | --- | --- | --- | --- |
|  | PLWH n (%) | PLw/oH n (%) | p-Value | Std. Diff | PLWH n (%) | PLw/oH n (%) | p-Value | Std. Diff |
| **Total n of patients** | 1087 (100%) | 394,437 (100%) |  |  | 1082 (100%) | 1082 (100%) |  |  |
| **Age at Index (years ± SD)** | 58.60±12.22 | 62.78±15.57 | 0.0000 | 0.3007 | 58.59±12.25 | 58.70±13.08 | 0.8347 | 0.0090 |
| **Sex** |  |  |  |  |  |  |  |  |
| Male | 877 (80.7%) | 204,376 (51.8%) | 0.0000 | 0.6408 | 872 (80.6%) | 876 (81.0%) | 0.8272 | 0.0094 |
| Female | 193 (17.8%) | 179,139 (45.4%) | 0.0000 | 0.6235 | 193 (17.9%) | 190 (17.6%) | 0.8658 | 0.0073 |
| Unknown Gender | 17 (1.6%) | 10,832 (2.8%) | 0.0171 | 0.0815 | 17 (1.6%) | 16 (1.5%) | 0.8608 | 0.0075 |
| **Ethnicity** |  |  |  |  |  |  |  |  |
| Hispanic or Latino | 45 (4.1%) | 5,908 (1.5%) | 0.0000 | 0.1601 | 44 (4.1%) | 55 (5.1%) | 0.2578 | 0.0487 |
| Not Hispanic or Latino | 827 (76.1%) | 279,817 (70.1%) | 0.0002 | 0.1163 | 823 (76.1%) | 822 (76.0%) | 0.9599 | 0.0022 |
| Unknown Ethnicity | 215 (19.8%) | 108,622 (27.6%) | 0.0000 | 0.1835 | 215 (19.9%) | 205 (19.0%) | 0.5868 | 0.0234 |
| **Race** |  |  |  |  |  |  |  |  |
| White | 916 (84.2%) | 326,200 (82.7%) | 0.1772 | 0.0418 | 915 (84.6%) | 924 (85.4%) | 0.5881 | 0.0233 |
| Black or African American | 78 (7.2%) | 4,104 (1.0%) | 0.0000 | 0.3129 | 74 (6.8%) | 72 (6.7%) | 0.8639 | 0.0074 |
| Asian | ≤10 | 3242 (0.8%) | 0.7213 | 0.0105 | ≤10 | ≤10 | 1.0000 | 0.0000 |
| Native Hawaiian or Other Pacific Islander | ≤10 | 1567 (0.4%) | 0.0063 | 0.0646 | ≤10 | ≤10 | 1.0000 | 0.0000 |
| American Indian or Alaska Native | ≤10 | 536 (0.1%) | 0.0000 | 0.1083 | ≤10 | 0 (0%) | 0.0015 | 0.1383 |
| Unknown Race | 70 (6.4%) | 48,985 (12.4%) | 0.0000 | 0.2058 | 70 (6.5%) | 65 (6.0%) | 0.6567 | 0.0191 |
| Other Race | 17 (1.6%) | 9713 (2.5%) | 0.0560 | 0.0640 | 17 (1.6%) | 16 (1.5%) | 0.8608 | 0.0075 |
| **Metastatic Sites** |  |  |  |  |  |  |  |  |
| Lymph Node | 52 (4.8%) | 5,101 (1.3%) | 0.0000 | 0.2044 | 51 (4.7%) | 51 (4.7%) | 1.0000 | 0.0000 |
| Lung | 14 (1.3%) | 2,169 (0.6%) | 0.0010 | 0.0774 | 14 (1.3%) | ≤10 | 0.4116 | 0.0353 |
| Brain / Meningeal | 21 (1.9%) | 2,116 (0.5%) | 0.0000 | 0.1266 | 21 (1.9%) | 18 (1.7%) | 0.6278 | 0.0208 |
| Liver | ≤10 | 1,909 (0.5%) | 0.0389 | 0.0522 | ≤10 | ≤10 | 1.0000 | 0.0000 |
| **Comorbidities** |  |  |  |  |  |  |  |  |
| Ischemic heart disease | 195 (17.9%) | 37,038 (9.4%) | 0.0000 | 0.2508 | 191 (17.7%) | 187 (17.3%) | 0.8208 | 0.0135 |
| Chronic viral hepatitis | 85 (7.8%) | 1,273 (0.3%) | 0.0000 | 0.3864 | 80 (7.4%) | 76 (7.0%) | 0.7395 | 0.0142 |
| Chronic lower respiratory disease | 242 (22.3%) | 36,569 (9.3%) | 0.0000 | 0.3622 | 238 (22.0%) | 232 (21.4%) | 0.7544 | 0.0135 |
| Chronic kidney disease | 116 (10.7%) | 19,621 (5.00%) | 0.0000 | 0.2123 | 114 (10.5%) | 120 (11.1%) | 0.6779 | 0.0179 |
| Hypertensive heart disease | 33 (3.0%) | 7,490 (1.9%) | 0.0062 | 0.0733 | 31 (2.9%) | 28 (2.6%) | 0.6921 | 0.0170 |
| Type 1 Diabetes Mellitus | 22(2.0%) | 3,984 (1.0%) | 0.0009 | 0.0830 | 22(2.0%) | 19 (1.8%) | 0.6362 | 0.0203 |

PLWH experienced a worsened long-term prognosis with a significantly lower ten-year survival rate (73.04% vs 77.95%, PLWH vs. PLw/oH, p=0.00628, **Fig. 1A, Table S7**), also seen in previous reports ^7^. The two cohorts had a similar median follow-up time of approximately 3.4 years (PLw/oH) and 4.2 years (PLWH, **Table S8**). Our analysis focused on patients diagnosed during the ART era (98.4% of patients had ART treatment records; patients diagnosed before 2005 were not available). Staging information and BRAF status were only available for <1% of the population, and no robust statistical assessment could be performed on these aspects. Nevertheless, given the impact of metastasis on cancer mortality, we assessed the occurrence of melanoma spreading to its most frequent metastatic sites. At presentation, the PLWH cohort displayed significantly increased occurrence of metastasis for all the analyzed sites (**Table 1**). We then assessed the risk of metastasis occurring after melanoma diagnosis. No significant differences were detected between the two groups of patients in terms of early metastasis (1 day to 6 months after diagnosis, **Table S9**). While there were no significant differences in lung, liver, or lymph node metastases, PLWH had a significantly higher risk of late brain metastases (>6 months after diagnosis, p=0.0009, risk ratio = 1.78, **Table S9**), and an overall increased risk of metastatic disease in the long term (p=0.0296, risk ratio = 1.325, **Table S9**). PLWH have also been reported to experience higher rates of chronic conditions affecting the heart, lungs, and liver, as well as diabetes, a trend that we observed in our cohort as well (**Table 1**). To account for potential confounders, including metastasis, other common chronic conditions, and demographic factors, we balanced the cohorts for these covariates (**Table 1**) and found significantly different survival rates even after adjustment (73.37% vs 81.16% survival rate, PLWH vs. PLw/oH, p=0.02411, **Fig. 1B, Table S10**).

**Figure 1.**
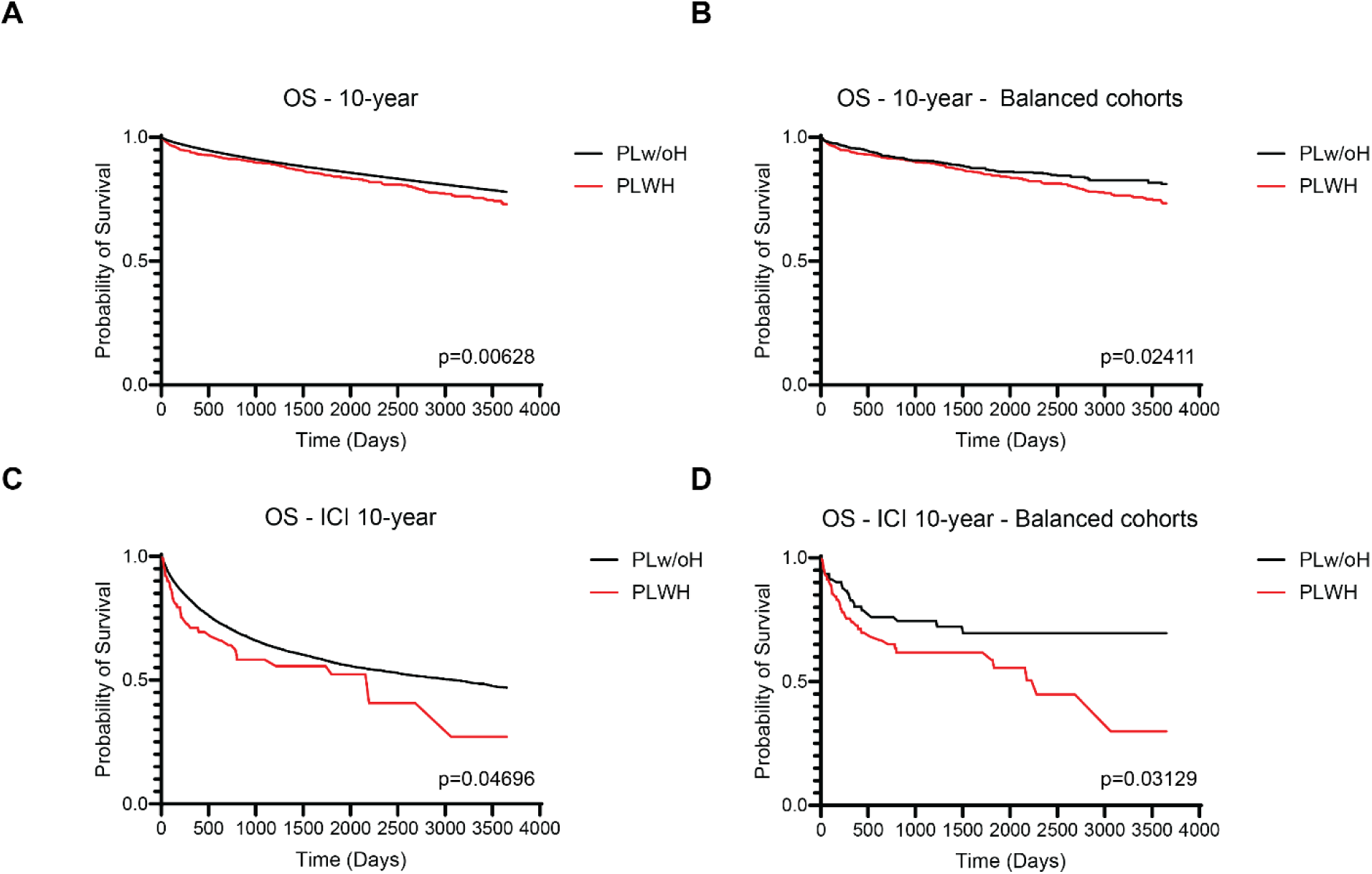
Overall survival rates and immune checkpoint therapy survival rates of PLWH and PLw/oH with melanoma. A-B) Kaplan-Meier graphs representing 10-year survival probability of PLWH and PLw/oH with melanoma. The index event is a melanoma diagnosis. A) Overall survival (OS), including all patients in the cohorts. B) OS after balancing for demographic covariates. C-D) Kaplan-Meier graphs representing 10-year survival probability of PLWH and PLw/oH with melanoma who underwent immune checkpoint therapy. The index event is the treatment start. C) OS including all patients in the cohorts. D) OS after balancing for demographic and clinical covariates. Reported p-values are from a Log-rank sum test between the two cohorts (p<0.05 is considered statistically significant).

Next, we focused on FDA-approved immune checkpoint inhibitors (ICIs) to evaluate differences in treatment patterns and outcomes between the two groups. ICIs serve as the first-line treatment approach for patients with advanced melanoma, and analyzing their use provides insights into how patients with late-stage disease fare in the clinical setting. Among our cohorts, approximately 9% of PLWH (n=95) and 6% of PLw/oH (n=23,827) had a record of receiving immune checkpoint therapies. The demographic features of these two sub-groups reflected most of the differences we detected between the parent cohorts (**Table 2**). Although we were not statistically powered enough to draw conclusions for specific treatment regimens, we were able to make some comprehensive observations regarding ICI outcomes and timing. We found that, on average, PLWH experienced longer delays before initiating treatment compared to PLw/oH (p=0.0026, **Tables S11-12**). In the PLWH cohort, we did observe a significantly higher number of patients with lymph node metastasis at presentation (**Table 2**). Once ICI was initiated, we did not detect differences in treatment duration (**Table S12**). We then performed a ten-year survival analysis of the two cohorts. PLWH demonstrated significantly lower probabilities of survival compared to the non-HIV melanoma patients (10-year, 27.14% vs 46.91% survival rate, PLWH vs. PLw/oH, p=0.04696, **Fig. 1C**, and **Table S13**). Such significant differences persisted even after adjusting for demographic covariates, metastasis at presentation, and major co-morbidities (10-year, 29.92% vs 69.54% survival rate, PLWH vs. PLw/oH, p=0.03129, **Fig. 1D, Table 1, and Table S14)**

**Table 2.** Demographics of PLWH and PLw/oH with melanoma cohorts for immunotherapy survival calculations, before and after cohort balancing.

|  | Before balancing cohorts |  |  |  | After balancing cohorts |  |  |  |
| --- | --- | --- | --- | --- | --- | --- | --- | --- |
|  | PLWH n (%) | PLw/oH n (%) | p-Value | Std. Diff | PLWH n (%) | PLw/oH n (%) | p-Value | Std. Diff |
| **Total n of patients** | 95 (100%) | 23,827 (100%) |  |  | 94 (100%) | 94 (100%) |  |  |
| **Age at Index (years ± SD)** | 61.91 ± 11.48 | 64.27 ± 14.58 | 0.1141 | 0.1804 | 61.15 ± 11.53 | 63.15 ± 12.65 | 0.4744 | 0.1046 |
| **Sex** |  |  |  |  |  |  |  |  |
| Male | 75 (79.0%) | 14,408 (60.5%) | 0.0002 | 0.4105 | 75 (79.8%) | 74 (78.7%) | 0.8572 | 0.0262 |
| Female | 18 (19.0%) | 8,878 (37.3%) | 0.0002 | 0.4161 | 17 (18.1%) | 17 (18.1%) | 1.0000 | 0.0000 |
| Unknown Gender | ≤10 | 541 (2.3%) | 0.0000 | 0.3422 | ≤10 | ≤10 | 1.0000 | 0.0000 |
| **Ethnicity** |  |  |  |  |  |  |  |  |
| Not Hispanic or Latino | 77 (81.1%) | 18,440 (77.4%) | 0.3944 | 0.0903 | 76 (80.9%) | 72 (76.0%) | 0.4760 | 0.1041 |
| Hispanic or Latino | ≤10 | 525 (2.2%) | 0.0000 | 0.3460 | ≤10 | ≤10 | 1.0000 | 0.0000 |
| Unknown Ethnicity | 15 (15.8%) | 4,862 (20.4%) | 0.2650 | 0.1201 | 15 (16.0%) | 19 (20.21%) | 0.4485 | 0.1107 |
| **Race** |  |  |  |  |  |  |  |  |
| White | 84 (88.4%) | 21,016 (88.2%) | 0.9474 | 0.0068 | 83 (88.3%) | 83 (88.3%) | 1.0000 | 0.0000 |
| Black or African American | ≤10 | 306 (1.3%) | 0.0000 | 0.3998 | ≤10 | ≤10 | 1.0000 | 0.0000 |
| Asian | ≤10 | 232 (1.0%) | 0.0000 | 0.4193 | ≤10 | ≤10 | 1.0000 | 0.0000 |
| Native Hawaiian / Pacific Islander | 0 (0%) | 23 (0.1%) | 0.7619 | 0.0440 | 0 (0%) | 0 (0%) |  |  |
| American Indian / Alaska Native | 0 (0%) | 34 (0.1%) | 0.7125 | 0.0534 | 0 (0%) | 0 (0%) |  |  |
| Other | ≤10 | 583 (2.5%) | 0.0000 | 0.3326 | ≤10 | ≤10 | 1.0000 | 0.0000 |
| Unknown | ≤10 | 1,633 (6.19%) | 0.1577 | 0.1307 | ≤10 | ≤10 | 1.0000 | 0.0000 |
| **Metastatic Sites** |  |  |  |  |  |  |  |  |
| Lymph Node | 45 (47.4%) | 8,082 (33.9%) | 0.0057 | 0.2764 | 44 (46.8%) | 44 (46.8%) | 1.0000 | 0.0000 |
| Lung | 12 (12.6%) | 3,259 (13.7%) | 0.7670 | 0.0309 | 12 (12.8%) | 11 (11.7%) | 0.8239 | 0.0325 |
| Brain / Meningeal | 13 (13.7%) | 2,364 (9.92%) | 0.2211 | 0.1168 | 13 (13.8%) | 12 (12.8%) | 0.8299 | 0.0313 |
| Liver | ≤10 | 1,969 (8.3%) | 0.4243 | 0.0776 | ≤10 | ≤10 | 1.0000 | 0.0000 |
| **Comorbidities** |  |  |  |  |  |  |  |  |
| Ischemic heart disease | 25 (26.3%) | 4,351 (18.3%) | 0.0427 | 0.1945 | 25 (26.6%) | 23 (24.5%) | 0.7380 | 0.0489 |
| Chronic viral hepatitis | ≤10 | 144 (0.6%) | 0.0000 | 0.4433 | ≤10 | ≤10 | 1.0000 | 0.0000 |
| Chronic lower respiratory disease | 24 (25.3%) | 3,898 (16.4%) | 0.0193 | 0.2207 | 23 (24.5%) | 28 (29.8%) | 0.4121 | 0.1198 |
| Chronic kidney disease | 18 (18.9%) | 2,069 (8.7%) | 0.0001 | 0.3008 | 17 (18.1%) | 18 (19.2%) | 0.8514 | 0.0273 |
| Hypertensive heart disease | ≤10 | 978 (4.1%) | 0.0017 | 0.2485 | ≤10 | ≤10 | 1.0000 | 0.0000 |
| Type 1 Diabetes Mellitus | ≤10 | 343 (1.4%) | 0.0000 | 0.3903 | ≤10 | ≤10 | 1.0000 | 0.0000 |

### PLWH display an immunosuppressive and exhausted melanoma tumor microenvironment

Our data indicate that PLWH who develop melanoma experience an overall elevated mortality rate, also in the sub-cohort who received ICI. While demographic differences, higher rates of coexisting conditions, and psychosocial barriers are contributors to mortality, the decreased survival rate persisted even after balancing cohorts for demographic and clinical covariates, indicating there may also be underlying biological or immunological components. To explore this, we collected tumor samples from 11 PLWH with melanoma and matched them with samples from 11 PLw/oH with melanoma with similar clinical and demographic features (**Table S1)**.

For each sample, three to five regions of interest (ROIs) were selected for spatial immune transcriptomic profiling (72 immune-related genes). A dermatopathologist defined ROIs based on morphological features and immunofluorescence staining for melanocytic markers, designating subregions as either tumor-cell–rich (TUMOR) or tumor-microenvironment-rich (TME) (**Fig. S1**). TUMOR ROIs corresponded to areas enriched in melanoma cells (positive for S100B/Pmel17) with limited immune infiltration, whereas TME ROIs encompassed CD45⁺ immune- and stroma-rich regions lacking tumor marker expression. This classification was designed to capture both tumor-invasive immune niches and the surrounding immune microenvironment. To identify transcriptional differences related to HIV status, we performed differential expression analysis (DEGs) using the DESeq2 algorithm, comparing both TME and TUMOR regions between PLWH and PLw/oH. The analysis revealed significant gene expression differences between the groups, highlighting distinct immune signatures and checkpoint expression patterns associated with HIV status (**Fig. 2**).

**Figure 2.**
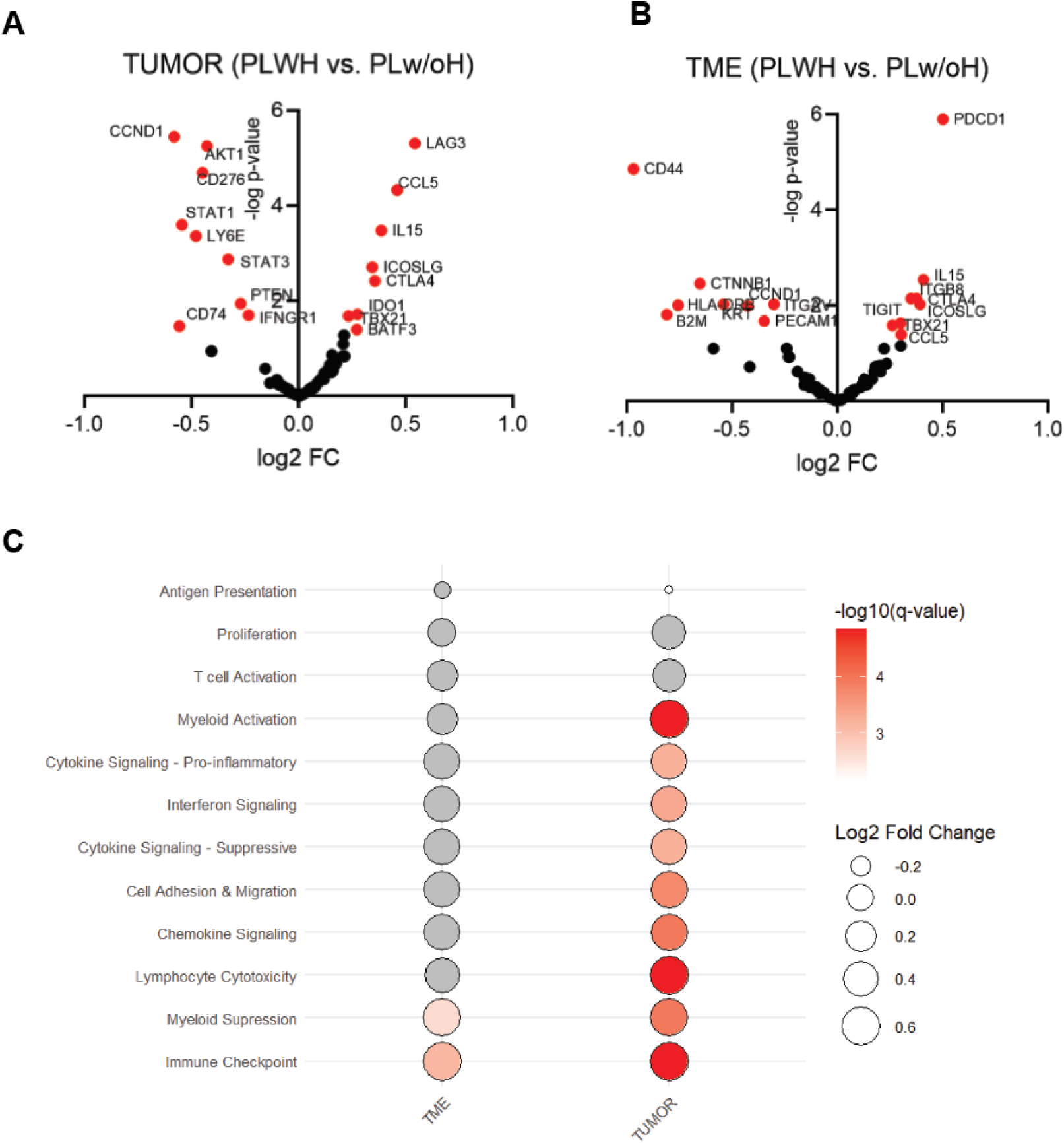
Spatial immune transcriptomic profiling of melanoma samples from PLWH and PLw/oH. Volcano plots for A) TME and B) TUMOR, comparing immune gene expression of PLWH vs PLw/oH melanoma samples (n=11); log2 fold change is on the x-axis, and -log10 of adjusted p-value is on the y-axis. Values were calculated by the DEseq2 algorithm (data points in red have an adjusted p-value <0.05). C) Bubble plot representation of immune signaling gene set analysis performed by running the MCPcounter algorithm on the gene expression matrix. Bubble plot size represents the log2 fold change of MCPcounter scores (PLWH vs. PLw/oH), and bubble color represents the adjusted p-value (q-value in a multiple Welch t-test with FDR of 0.01) for the TUMOR and TME areas (PLWH vs. PLw/oH). A q-value<0.01 was considered significant; comparisons with non-significant q-values are represented in grey.

When comparing the TME and TUMOR areas of PLWH with PLw/oH, we observed increased expression of several immune checkpoint genes, including *PD1*, *LAG3*, *CTLA4*, *TIGIT*, *CD276,* and *ICOSLG*. Notably, *LAG3* emerged as the top-scoring immune checkpoint in the TUMOR, while *PD1* was the highest in the TME (**Fig. 2A- B**). This suggests that there are distinct spatial distributions of varying T cell populations within these tumors. Both the TUMOR and TME exhibited additional characteristics of an immune-suppressive or evasive microenvironment. In the TUMOR compartment, we observed elevated expression of *IDO1*, a key protein that inhibits T-cell function (**Fig. 2A**). Meanwhile, the TME showed reduced expression of *HLA-DRB* and *B2M,* which are MHC class II and I molecules essential for antigen presentation (**Fig. 2B**). Several key genes involved in immune responses were also downregulated, including *IFNGR1*, *CD74*, *LY6E*, and *STAT2*. Beyond immune regulation, we identified a pattern of differential gene expression pertaining to fibrosis and extracellular matrix (ECM) remodeling in the TME. PLWH demonstrated a decline in the expression of *CD44*, *ITGAV*, *and PECAM1* and increased *ITGB8*, suggesting alterations in tumor-stroma interactions that may contribute to disease progression (**Fig. 2B**).

Additionally, a small cadre of genes associated with enhanced inflammation was identified in the TME of PLWH, including an increase in *IL-15*, *TBX21*, and *CCL5* (**Fig. 2B**). In the TUMOR, there was an increase in *BATF3*, which is mostly expressed on CD8^+^ cells (i.e., dendritic cells and T cells) (**Fig. 2A**).

To summarize our findings and prioritize downstream validation, we ran a modified version of the MCPcounter algorithm, utilizing tailored gene sets that included population marker genes and immune signaling genes (**Tables S3-S4**). The only two upregulated gene sets in PLWH, both in TME and TUMOR areas, were immune checkpoints and myeloid suppressive signaling (**Fig. 2C**). In terms of immune populations, overall, it appeared that the TUMOR areas from PLWH were generally enriched in immune cells both from myeloid and lymphoid origin and displayed enhanced expression of several immune signaling pathways (**Fig. 2C and Fig. S2**). The TME areas were, in general, more similar between PLWH and PLw/oH (**Fig. S2**).

### PLWH display distinct subsets of CD8^+^ T cells within the melanoma tumor environment

To validate and further expand upon the spatial transcriptomics findings, we investigated T cell populations using multiplexed immunofluorescence staining (mIF). The cohort included melanoma tumor samples from 15 HIV^+^ and 14 HIV^-^ patient samples, with 11 samples in each group overlapping with those profiled by spatial transcriptomics (**Fig. S3** and **Table S1**). We examined T cell populations within both the tumor bulk and peripheral edges using an mIF panel targeting T cell populations (CD8, CD4, Foxp3), along with the two top-scoring immune checkpoint inhibitors identified in our transcriptional analysis (LAG3 and PD1) (**Fig 3A**). While melanocytic markers were excluded from this panel, a pathologist histologically assessed the sections to delineate tumor regions and guided the analysis. Thus, the results reflect the tumor specimens as a whole instead of the specific delineation of sub-regions.

**Figure 3.**
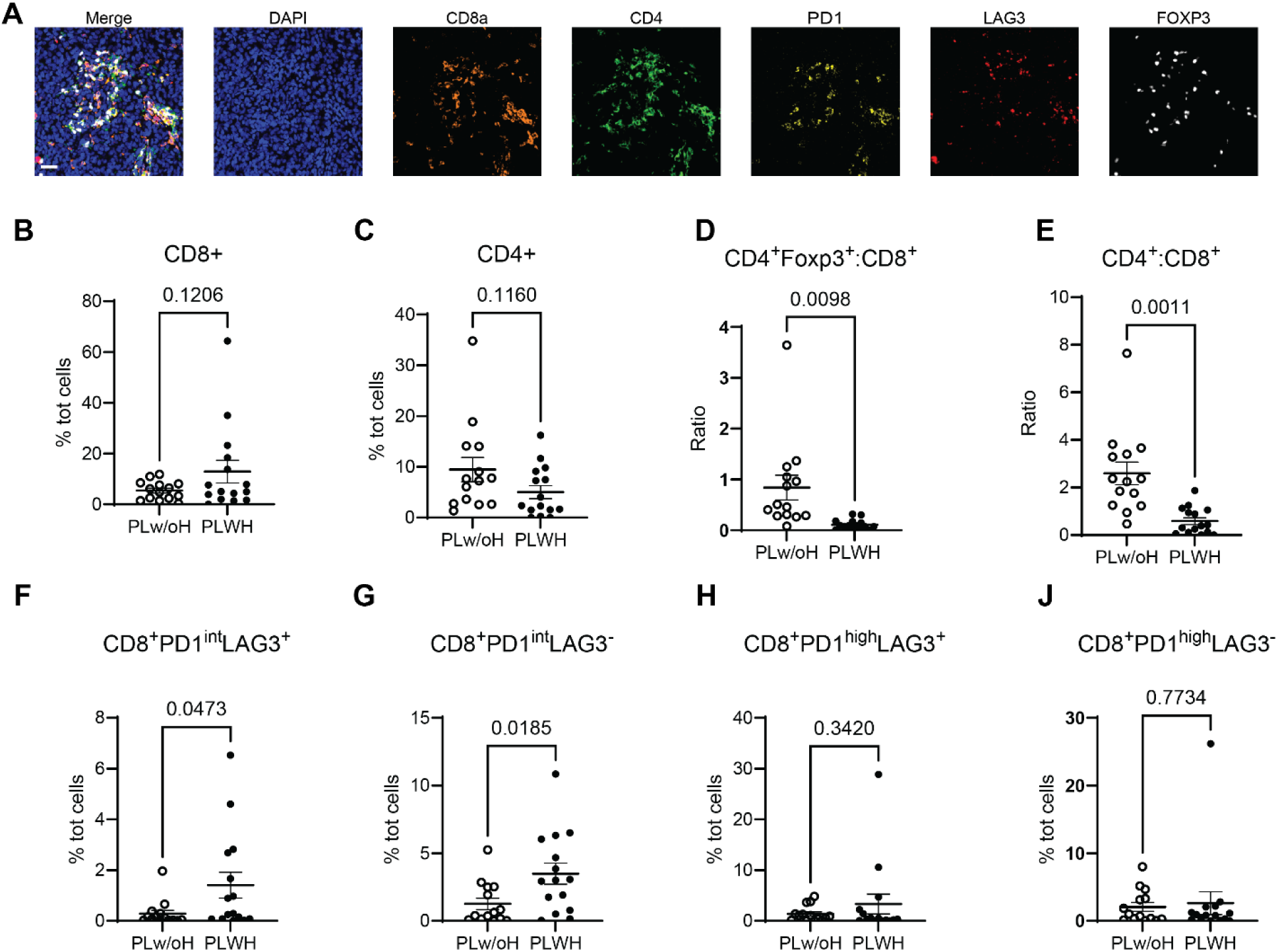
Multiplex IF assessment of T-cell populations’ abundance and immune checkpoint expression in melanoma samples from PLWH and PLw/oH. A) Representative images depicting each staining marker of our T cell multiplex immunofluorescence (mIF) panel, which was used to analyze melanoma tumors from PLWH and PLw/oH. The scale bar represents 50 µm. Quantification of mIF was performed on PLWH (n=15) and PLw/oH (n=14) tumor samples. Percentage of B) CD8^+^ and C) CD4^+^ T cells over the total number of cells. D) Ratio of CD4^+^Foxp3^+^ to CD8^+^ T cells, and E) CD4^+^ T cells to CD8^+^ T cells. Percentage of PD1^int^ CD8^+^ T cells that are F) LAG3^+^ and G)LAG3^-^ over the total number of cells. Percentage of PD1^HIGH^ CD8^+^ T-cells that are H) LAG3^+^ and J) LAG3^-^ over the total number of cells. The represented p-values are from a Welch’s t-test; p < 0.05 is considered significant, and error bars represent SEM.

Analysis of the T cell compartment revealed no significant differences in the overall frequency of CD8^+^ or CD4^+^ T cells between melanoma tumors in PLWH and uninfected patients (**Fig. 3B-C**). Similarly, regulatory T cells (T_REG_, CD4^+^FOXP3^+^) abundance did not differ significantly between groups (**Fig. S4A**). Notably, PLWH exhibited altered T cell proportions, with a significantly reduced CD4^+^:CD8^+^ ratio and a lower Foxp3^+^:CD8^+^ ratio, suggesting a shift in the relative balance of T cell subsets (**Fig. 3D-E**).

We next interrogated the co-expression of immune checkpoint receptors PD1 and LAG3 on CD4^+^ and CD8^+^ T cells. PD1 expression was stratified into three tiers (PD1^-^, PD1^int^, and PD1^high^), and LAG3 into two (LAG3^-^ and LAG3^+^), based on previous reports highlighting functional differences in cytotoxicity and exhaustion in these subpopulations^12–14^. No significant differences were observed in CD4^+^ T cell subsets between PLWH and PLw/oH (**Fig. S4B-E**).

Notably, we identified distinct subpopulations of CD8^+^ T cells in PLWH compared with PLw/oH. Specifically, melanoma tumors from PLWH showed significant enrichment of the CD8^+^PD1^int^LAG3^-^ and the CD8^+^PD1^int^LAG3^+^ populations (**Fig. 3F-G**). Notably, we did not detect any differences in the abundance of PD1^HIGH^ cells, whether they were LAG3^+^ or LAG3^-^ (**Fig. 3H-J**).

### PLWH exhibit an enriched population of immunosuppressive myeloid cells in melanoma tumors

Given the important role of the myeloid compartment in modulating T cell function, we next turned our focus to tumor-associated myeloid populations. Pathway analysis highlighted myeloid suppressive signaling as a top candidate (**Fig. 2C**) and a significant downregulation of key components of antigen presentation, such as *HLA-DRB*, within the tumor regions (**Fig. 2**). In parallel, previous studies have reported that PLWH have elevated levels of myeloid-derived suppressor cells (MDSCs) at baseline, a population of highly immunosuppressive, immature myeloid cells ^15,16^. Based on these findings, we selected CD11b to broadly define the myeloid lineage and included HLA-DR, CD33, and CD66b to resolve subpopulations (**Fig. 4A**), including potential MDSCs. Using multiplexed immunofluorescence (mIF) on the tumor samples previously analyzed for T cell populations, we identified a significant increase in CD11b⁺ HLA-DR⁻ myeloid cells in PLWH (**Fig. 4B**), indicating an immature or non-antigen-presenting phenotype. Most of these cells also expressed CD33, supporting their classification as monocytic-like MDSCs (M-MDSCs) (**Fig. 4C**). In contrast, no differences were observed in CD11b⁺ HLA-DR⁻ CD66b⁺ cells, or polymorphonuclear-like MDSCs (PMN-MDSC) between PLWH and PLw/oH (**Fig. 4D**) ^17^.

**Figure 4.**
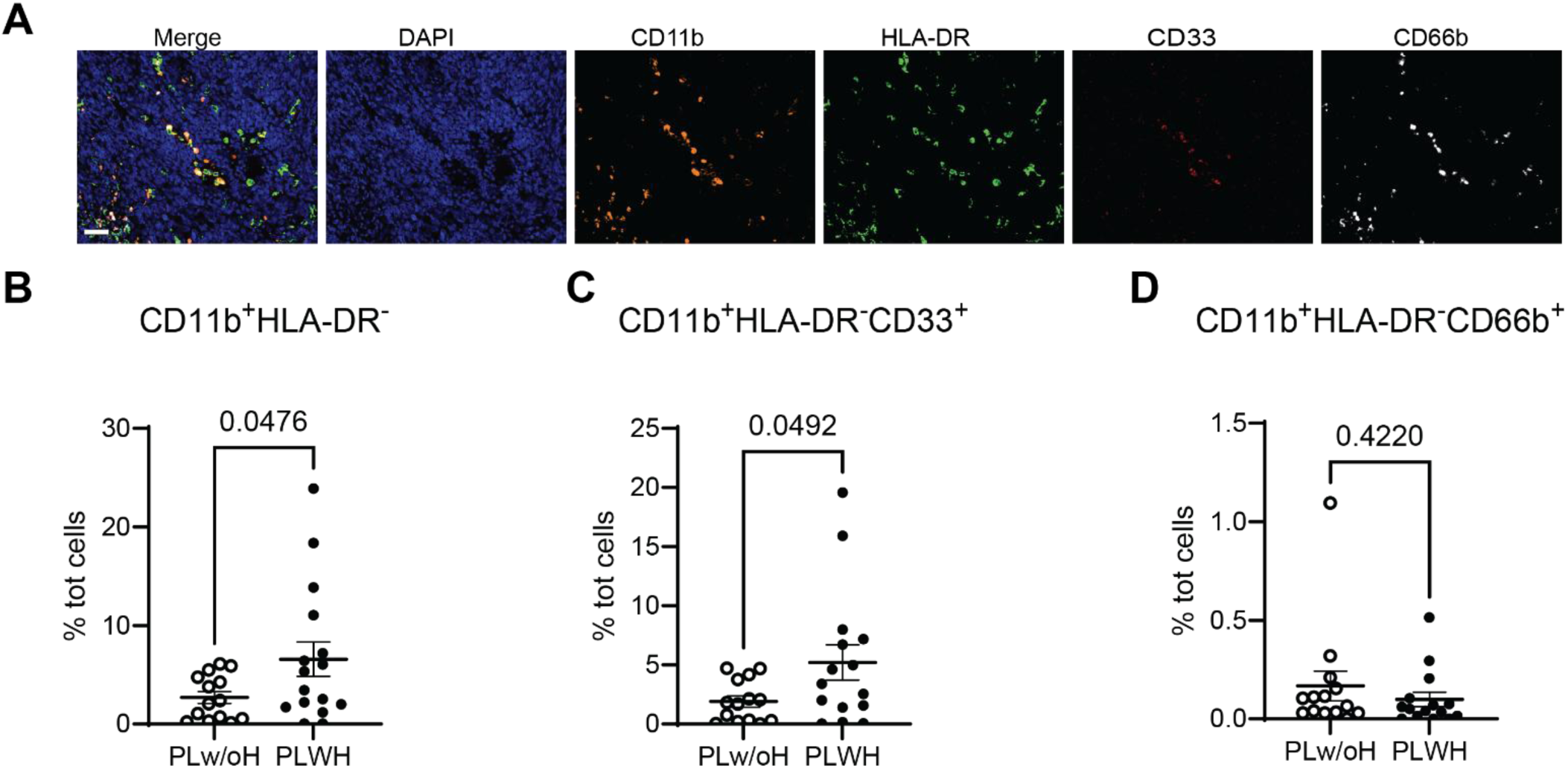
Multiplex IF assessment of myeloid populations’ abundance in melanoma samples from PLWH and PLw/oH. A) Representative images depicting each staining marker used in our myeloid multiplex immunofluorescence (mIF) panel, used to analyze melanoma tumor samples from PLWH and PLw/oH. The scale bar represents 50 µm. Quantification of mIF was performed on PLWH (n=15) and PLw/oH (n=14) tumor samples. Percentage of B) CD11b^+^HLA-DR^-^, C) CD11b^+^HLA-DR^-^, CD33^+^, D) CD11b^+^HLA-DR^-^, CD66b^+^, over the total number of cells. The represented p-values are from a Welch’s t-test; p < 0.05 is considered significant, and error bars represent SEM.

## Discussion

The present study demonstrates that PLWH who develop melanoma experience distinct clinical outcomes and tumor microenvironments compared to those without HIV. Notably, PLWH exhibited increased overall mortality, including in the subpopulation of patients who underwent ICI, even after adjusting for demographic and clinical covariates. Immune-focused spatial transcriptomics analysis revealed a more suppressive and likely exhausted tumor microenvironment (TME) in PLWH, which was further supported by multiplex immunofluorescence identifying unique CD8^+^ T cell subsets and an increased presence of myeloid derived suppressor cells (MDSCs).

Despite increasing evidence of ICI safety in PLWH^18^, this patient population has been largely excluded from oncology clinical trials, resulting in limited efficacy data and persistent provider hesitancy ^19,20^. This exclusion reflects broader patterns of delayed cancer treatment in PLWH, potentially driven by disparities in healthcare access, treatment prioritization, and clinical decision-making. Our findings echo these systemic gaps, demonstrating that PLWH experience significant delays in treatment initiation, likely driven by both clinical uncertainty and psychosocial barriers, including limited access to care and persistent medical mistrust rooted in historical HIV-related stigma, as we have previously reviewed ^21^. Addressing these barriers is critical to improving melanoma outcomes and ensuring equitable cancer care for PLWH.

We also found a significantly higher long-term risk of metastasis in PLWH, particularly to the brain. As HIV infection is well known to develop into HIV-associated neurocognitive disorder (HAND)^22^, this may reflect an HIV-related disruption of the blood-brain barrier, creating a permissive environment for melanoma colonization^23^. Given that brain metastasis significantly lowers survival and treatment responses, this highlights the importance of heightened cancer surveillance and early treatment strategies for this population^24,25^.

In the ICI-treated cohort, none of whom received prior targeted therapy - an established factor for ICI responsiveness - PLWH displayed decreased survival despite similar time on treatment. These data raise the possibility that ICI may not confer the same durable responses in PLWH, which warrants further investigation^26^.

In addition to the clinical characteristics of melanoma in PLWH, our study provides critical insights into how chronic HIV infection fundamentally alters the tumor immune landscape, potentially driving worsened outcomes for this population of patients. We identified two enriched CD8^+^ T cell subsets: CD8^+^ PD1^INT^ LAG3^+^ and CD8^+^ PD1^INT^ LAG3^-^. These subsets likely represent recently activated or pre-exhausted phenotypes, with LAG3 co-expression acting as a stronger predictor of exhaustion.

PD1 and LAG3 are inhibitory receptors upregulated on CD8⁺ T cells during chronic viral infections and within tumors. While initially markers of activation, sustained co-expression drives T cell exhaustion. Similar levels of CD8⁺ PD1^HIGH^ T cells observed in both PLWH and PLw/oH likely reflect chronic stimulation by tumor antigens or PD-L1 expression ^27–29^. However, PLWH showed increased CD8⁺ PD1^INT^ LAG3⁺/⁻ T cells, suggesting a population of recently activated or pre-exhausted T cells. LAG3 co-expression indicates progression towards exhaustion, with their fate influenced by tumor-derived cues. Supporting this interpretation, spatial transcriptomic analysis revealed elevated *LAG3* transcription localized near tumor cells, consistent with a spatially driven gradient of exhaustion. PD1 and LAG3 co-expression on CD8⁺ T cells has been linked to impaired anti-tumor immunity, and while dual blockade of PD1 and LAG3 has shown promise in improving progression-free survival in metastatic melanoma ^30–32^, clinical data on ICI efficacy in PLWH remain scarce, particularly for anti-LAG3 therapies. Further research is needed to define how these pathways shape tumor progression in PLWH and inform targeted therapies to restore immune function.

Despite HIV’s well-documented impact on the CD4^+^ T cell compartment ^33,34^, we observed no significant differences in total CD4^+^ T cell counts or their subsets between cohorts, suggesting that immune dysregulation in the TME of PLWH is primarily driven by CD8^+^ T cells. This is further supported by a decreased CD4:CD8 ratio in PLWH.

Transcriptomic and multiplex IF analyses pointed to a shift in the myeloid compartment: a downregulation of antigen presentation genes and a shift from classical to cross-presentation mechanisms, indicative of an immune-evasive or immunosuppressive myeloid phenotype ^35–37^, supported by an accumulation of CD11b^+^ HLA-DR^-^ CD33^+^ cells in melanoma tumors of PLWH. These cells, defined as myeloid-derived suppressor cells (MDSCs), are a rare and potent immunosuppressive subset of immature myeloid cells. In healthy individuals, MDSCs typically emerge later in cancer progression and are known contributors to immune evasion and resistance to immunotherapy ^38,39^. However, PLWH exhibit elevated baseline levels of circulating MDSCs, regardless of ART status, likely due to chronic low-level inflammation^15,17^. This may promote early recruitment of MDSCs to the melanoma TME, accelerating melanoma progression in this population by impairing the T cell response ^15–17,40–42^. Our data support this hypothesis, as we observed an upregulation in the expression of *CCL5* in both TUMOR and TME regions. While CCL5 is best known for its role in pro-inflammatory T cell generation and its involvement in HIV pathogenesis, via CCR5 binding^43–46^, it also recruits MDSCs and has been implicated in immune suppression in melanoma^47–49^. Elevated MDSC levels prior to initiating immunotherapy in melanoma patients have been associated with poorer treatment response and reduced overall survival, and CD11b^+^ HLA-DR^-^ CD33^+^ cells have emerged as a predictive biomarker for advanced-stage melanoma prognosis ^50^. Collectively, our findings suggest that MDSCs play a central role in shaping the immune landscape of melanoma in PLWH, likely impairing CD8^+^ T cell responses and contributing to worse clinical outcomes.

Despite inclusion of all confidently identified samples with corresponding clinical data, this study is limited by incomplete information regarding melanoma subtypes and staging, and by the modest sample sizes available for transcriptomic and mIF analyses. Moreover, the absence of an independent validation cohort limits the generalizability of the results and the robustness of the conclusions. These limitations underscore the broader lack of well-annotated biospecimens in this population. Future studies integrating tumor stage, subtype, and demographic data will enable more comprehensive and interpretable analyses. Nonetheless, the significant differences identified here provide a strong rationale for future, more focused biological investigations. These should aim to isolate the immune features of primary versus metastatic tumors in PLWH, further characterize the immature phenotype and functional role of MDSCs in PLWH, and perform functional assays on CD8^+^ T cells to better understand how their altered states affect anti-tumor responses. Such efforts will be critical to identifying actionable targets within both the T cell and myeloid compartments, ultimately guiding strategies to restore T cell function or limit MDSC accumulation.

Our investigation demonstrates that, despite effective ART, HIV infection fundamentally alters melanoma progression, as evidenced by changes in CD8^+^ T cell subsets and an immunosuppressive myeloid cell presence within the tumors. These immune disruptions, combined with delayed healthcare responses and an increased risk for metastasis, likely contribute to the elevated mortality rate observed in PLWH. Collectively, these findings underscore the urgent need for inclusive clinical trials and targeted therapeutic strategies to address the unique challenges faced by PLWH and improve cancer outcomes for this population.

## Acknowledgments

We thank the AIDS and Cancer Specimen Resource (ACSR) for providing tumor samples from PLWH; ACSR is funded by the National Cancer Institute.

## Notes

Funding: G.R. is supported by: WW Smith Charitable Trust - Medical Research Program (#C2303); Department of Defense - Congressionally Directed Medical Research Programs, Melanoma Research Program (ME220048); American Cancer Society (DBG-23-1036360-01); Comprehensive NeuroHIV Center (CNHC) Developmental Research and Mentoring Core (NIMH - P30MH092177); Sidney Kimmel Comprehensive Cancer Center (#00023568 and #901333). L.B. is supported by the Translational Research Training Grant in NeuroHIV (NIMH, T32-MH079785). P.J.G is supported by the National Institute of Drug Abuse, DA057337 and DA058051. Experimental Pathology Research Laboratory [RRID: SCR_017928] is partially supported by the Cancer Center Support Grant (P30CA016087). The original Akoya/PerkinElmer Vectra® multispectral imaging system was awarded through the Shared Instrumentation Grant S10 OD021747.

Conflict of interest statement: The authors declare no conflicts of interest.

### Competing Interest Statement

The authors have declared no competing interest.

### Summary of Updates

Title was revised, new co-variate analysis was performed

