## Supplementary Figures for "Population analysis and immunologic landscape of melanoma in people living with HIV indicate impaired antitumor immunity"

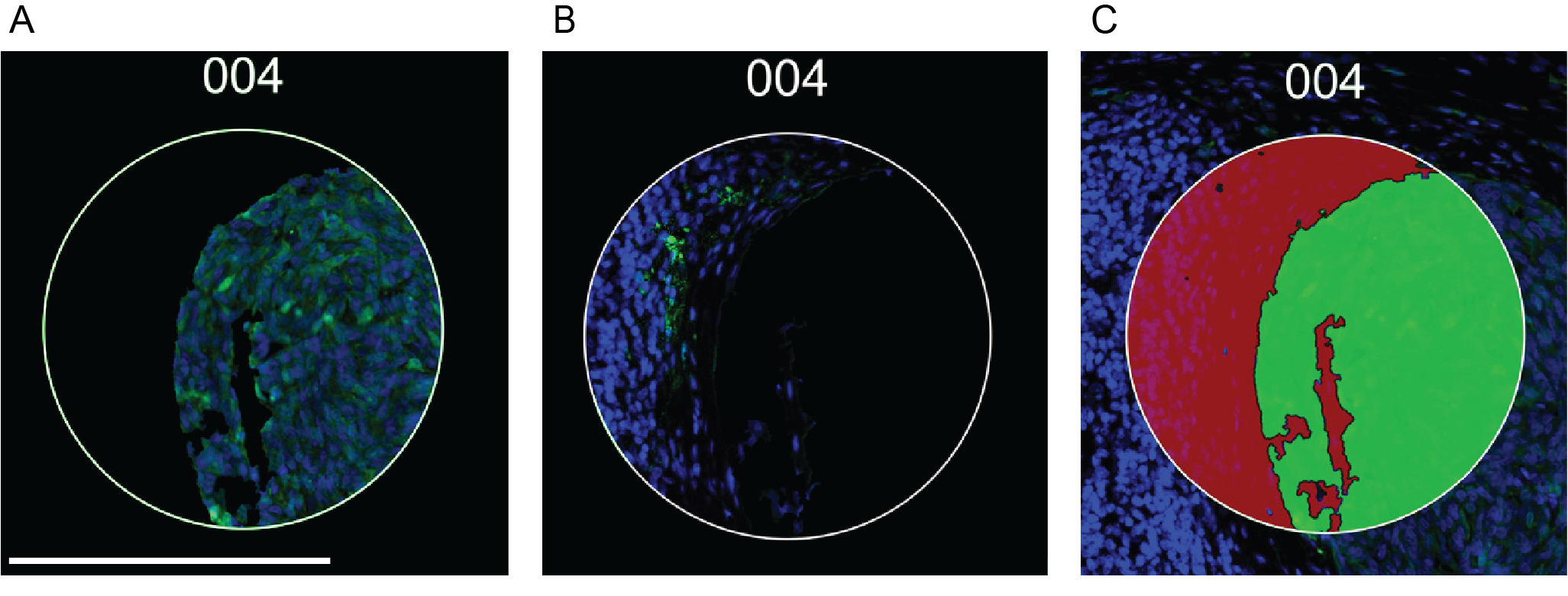

**Figure S1. Masking strategy for immune spatial transcriptomics.** Representative images illustrating the masking strategy used to distinguish tumor (TUMOR) and tumor microenvironment (TME) regions. For each sample, three to five regions of interest (ROIs) were selected for spatial immune transcriptomic profiling (72 immune-related genes). A dermatopathologist identified ROIs based on morphological features and immunofluorescence staining patterns. A) Tumor-rich (TUMOR) ROIs were characterized by cohesive nests of large cells with uniform nuclear size and shape and by positive staining for melanocytic markers S100B/Pmel17, with minimal infiltration by CD45⁺ immune cells (S100B/Pmel17, green; DAPI nuclear staining blue). B) Tumor-microenvironment-rich (TME) ROIs were selected based on the presence of CD45⁺ immune cells and absence of tumor marker expression (S100B/Pmel17⁻), in addition to distinct stromal or lymphocytic morphology (CD45, green; DAPI nuclear staining blue). C) Masks were applied according to this classification, with green masks corresponding to TUMOR and red masks corresponding to TME regions. Bar represents 250 μm.

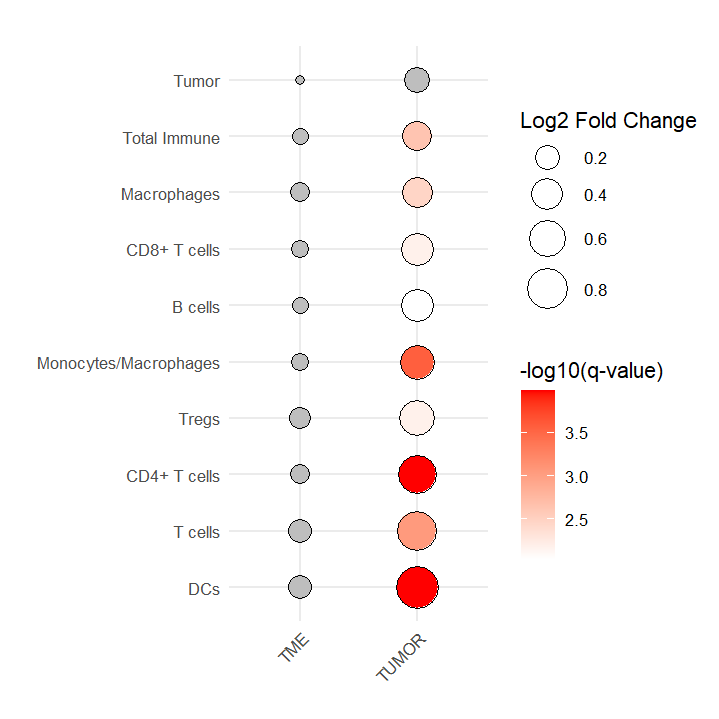

**Figure S2. Immune cell population deconvolution analysis on melanoma samples from PLWH and PLw/oH.** Bubble plot representation of marker gene set analysis performed using the MCPcounter algorithm on the immune gene expression matrix. Bubble plot size represents the log2 fold change of MCPcounter scores (PLWH vs. PLw/oH), and bubble color represents the adjusted p-value (q-value in a multiple Welch t-test with FDR of 0.0001) for the TUMOR and TME areas. A q-value<0.0001 was considered significant; comparisons with non-significant q-values are represented in grey.

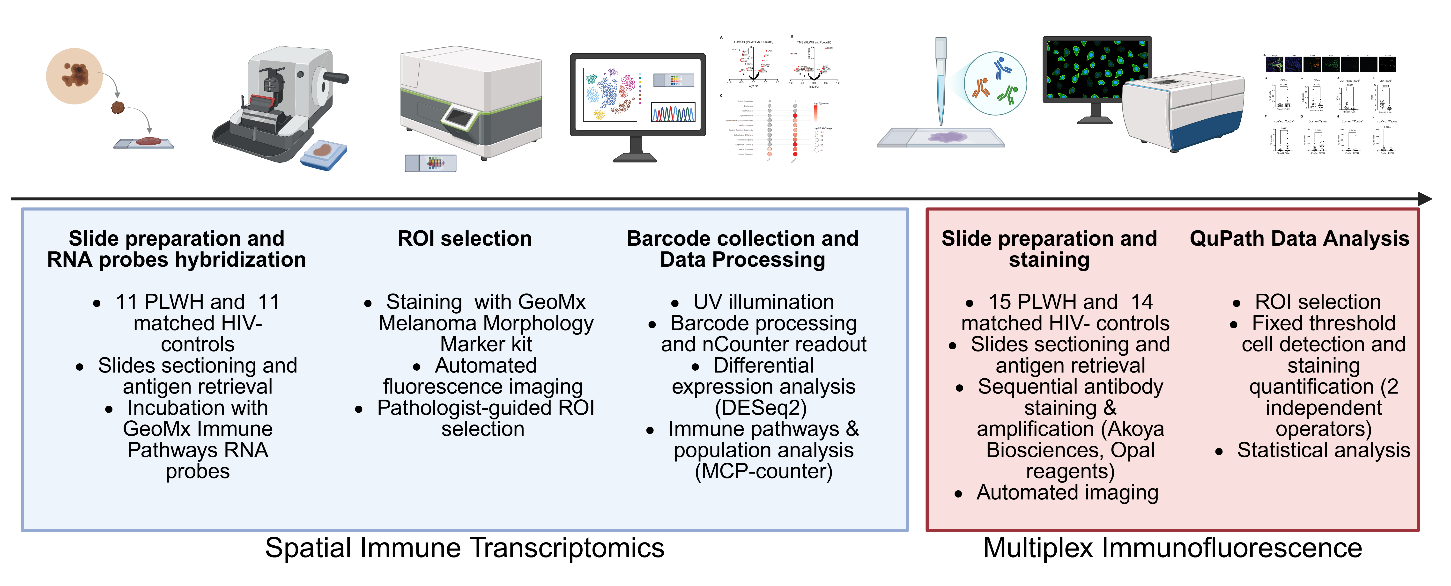

**Figure S3. Schematic representation of patient samples, assays, and analysis workflow.** The diagram indicates the major steps involved in tissue processing, spatial transcriptomic assay and analysis, and multiplex immunofluorescence assay and analysis. Created in BioRender. Romano, G (2025). <https://BioRender.com/kb979ps> .

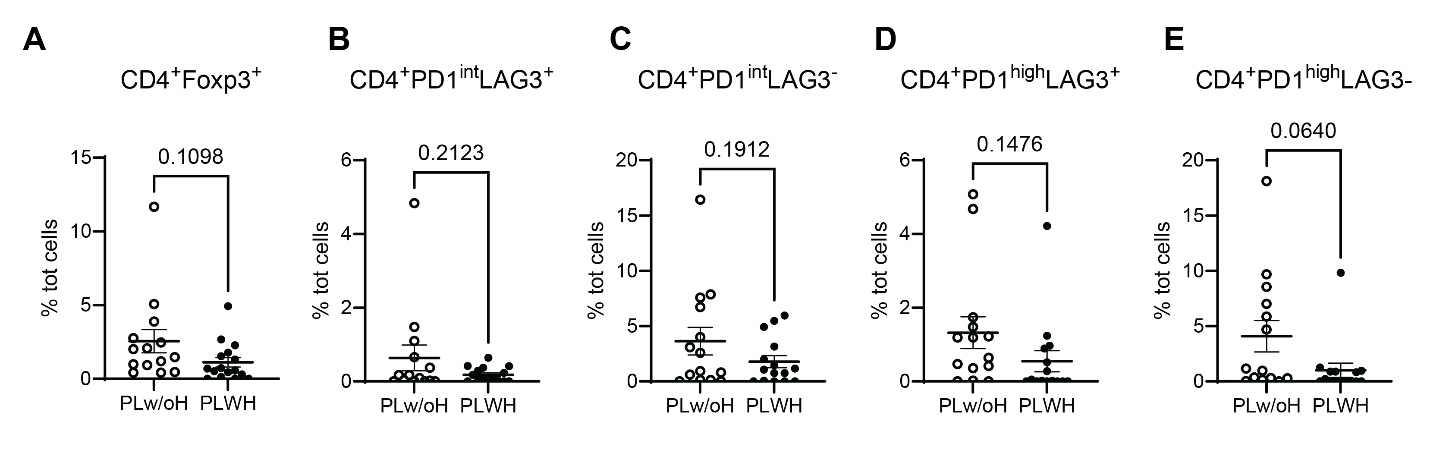

**Figure S4. Multiplex IF assessment of CD4+ T cell populations abundance and immune checkpoint expression in melanoma samples from PLWH and PLw/oH.**  Quantification of mIF was performed on PLWH (n=15) and PLw/oH (n=14) tumor samples. Percentage of A) CD4+Foxp3+ over the total number of cells. Percentage of PD1^int^ CD4+ T-cells that are B) LAG3+ and C) LAG3- over the total number of cells. Percentage of PD1^high^ CD4+ T-cells that are D) LAG3+ and E) LAG3- over the total number of cells. The represented p-values are from a Welch’s t-test; p < 0.05 is considered significant.
